# Discriminating between sleep and exercise-induced fatigue using computer vision and behavioral genetics

**DOI:** 10.1101/2020.02.06.937359

**Authors:** Kelsey N. Schuch, Lakshmi Narasimhan Govindarajan, Yuliang Guo, Saba N. Baskoylu, Sarah Kim, Benjamin Kimia, Thomas Serre, Anne C. Hart

## Abstract

Following prolonged swimming, *Caenorhabditis elegans* cycle between active swimming bouts and inactive quiescent bouts. Swimming is exercise for *C. elegans* and here we suggest that inactive bouts are a recovery state akin to fatigue. Previously, analysis of exercise-induced quiescent (EIQ) bouts relied on laborious manual observation, as existing automated analysis methods for *C. elegans* swimming either cannot analyze EIQ bouts or fail to accurately track animal posture during these bouts. It is known that cGMP-dependent kinase (PKG) activity plays a conserved role in sleep, rest, and arousal. Using *C. elegans* EGL-4 PKG, we first validate a novel learning-based computer vision approach to automatically analyze *C. elegans* locomotory behavior and distinguish between activity and inactivity during swimming for long periods of time. We find that *C. elegans* EGL-4 PKG function predicts EIQ first bout timing, fractional quiescence, bout number, and bout duration, suggesting that previously described pathways are engaged during EIQ bouts. However, EIQ bouts are likely not sleep as animals are feeding during the majority of EIQ bouts. We find that genetic perturbation of neurons required for other *C. elegans* sleep states also does not alter EIQ dynamics. Additionally, we find that EIQ onset is sensitive to age and DAF-16 FOXO function. In summary, we have validated a new behavioral analysis software that enabled a quantitative and detailed assessment of swimming behavior, including EIQ. We found novel EIQ defects in aged animals and animals with mutations in a gene involved in stress tolerance. We anticipate that further use of this software will facilitate the analysis of genes and pathways critical for fatigue and other *C. elegans* behaviors.

## Introduction

Fatigue is a commonly experienced phenomenon that is typically defined by a feeling of exhaustion combined with decreased muscle output. Feelings of fatigue are common after vigorous physical activity or exercise, but fatigue is also a hallmark symptom of a variety of health disorders and diseases, including cancer, mood disorders, neurodegenerative disorders, and chronic fatigue syndrome. Fatigue is not limited to vertebrates; exercise eventually drives decreased spontaneous locomotion in invertebrates as well. The molecular pathways and mechanisms involved in fatigue have not been fully delineated in any animal species.

The nematode *Caenorhabditis elegans* provides a potentially powerful model system to interrogate genetic mechanisms and cellular pathways underlying fatigue. Several methods have been developed to test the neuromuscular output of *C. elegans* during locomotion, including burrowing assays through media of varying densities (Beron *et al*., 2015) and pillar deflection strength measuring assays (Rahman *et al*., 2018). *C. elegans* are typically grown on solid media, but swimming exercise in liquid media has been shown to be energetically costly (Laranjeiro, Harinath, Burke, Braeckman, & Driscoll, 2017). Following prolonged swimming, *C. elegans* begin to spontaneously cycle between periods of active swimming with vigorous body undulations (active bouts) and periods of immobility that lack body undulations (quiescent bouts) (Ghosh & Emmons, 2008). Because swimming is exercise for *C. elegans*, these quiescent bouts may be fatigue, as they occur after the exertion of swimming and represent a decline in muscle output. Definition and dissection of exercise-induced quiescence (EIQ) pave the way to use this system to study conserved mechanisms fundamental to fatigue in all animals.

However, using *C. elegans* to study EIQ and fatigue requires analysis of swimming behavior over long periods of time, and therefore presents logistical and computational challenges. Historically, swimming behavior has been either 1) manually annotated at an extreme time cost on researchers or 2) sparsely sampled, which limited analysis depth. Several methods have been specifically developed for automatically estimating the pose of small laboratory animals, including *C. elegans* (Gomez-Marin, Partoune, Stephens, Louis, & Brembs, 2012; Jung, Aleman-Meza, Riepe, & Zhong, 2014; Patel *et al*., 2014; Restif *et al*., 2014). For the most part, methods developed for automatically estimating the pose of small laboratory animals rely on simple image processing (e.g., background subtraction) to extract the silhouette of a body before computing a medial axis transform. A major drawback of such methods is their inability to discriminate between the front and rear ends of the body, forcing researchers to rely on simple heuristics instead (Jung *et al*., 2014; Restif *et al*., 2014) (e.g., by computing the direction of movement and assuming that the animal moves forward). In addition, background subtraction tends to be sensitive to changes in illumination and often yields erroneous pose estimates. In the context of biological research, these failures need to be detected – either automatically (Restif *et al*., 2014) or manually (Jung *et al*., 2014; Stephens, Johnson-Kerner, Bialek, & Ryu, 2008) in order to exclude the corresponding frames from further behavioral analysis. Such an opportunistic approach to pose tracking may lead to significant biases in behavioral analyses if those system failures tend to co-occur more frequently with certain behaviors (e.g., for those behaviors that yield significant self-occlusion such as omega turns). Overall, existing approaches are not robust enough to allow for the throughput needed for behavioral analysis in modern biological research applications.

Here, we extended previous computer-vision work (Yang & Ramanan, 2013) and developed a learning-based approach (see Methods and (Guo, Govindarajan, Kimia, & Serre, 2018) for details) which outperforms other approaches including a deep neural network-based approach that exhibits state-of-the-art accuracy for human tracking (unlike humans, *C. elegans* lack distinctive body parts presenting a significant challenge for deep neural networks and related appearance-based approaches). The resulting system is able to efficiently distinguish between periods of activity and inactivity in freely swimming *C. elegans* (Guo *et al*., 2018) – addressing the need for high-resolution analysis that can handle both extended periods of swimming and quiescent behaviors in *C. elegans*. This system also permits fine-grained analysis of body movements and assessment of locomotion changes in *C. elegans* swimming over time and additionally handles complex postures more accurately than previously described systems.

Here, we use this new computer vision system to examine *C. elegans* locomotory behaviors and EIQ after prolonged swimming. We validate this system using mutant strains known to have altered EIQ (Ghosh & Emmons, 2008) and determine that most EIQ bouts are not a sleep state, as animals continue pharyngeal pumping during EIQ bouts. Further, we describe how EIQ changes over time with extended free swimming and report previously undescribed changes in EIQ as animal age. This new computer-vision analysis system should be valuable for *C. elegans* researchers in any field that requires accurate assessment of locomotory behavior over extended time intervals.

## Methods

### Strains and maintenance

Wild type N2 Bristol, MT1072 *egl-4(n477)*, DA521 *egl-4(ad450)*, HBR227 *aptf-1(gk794)*, HBR232 *aptf-1(tm3287)*, IB16 *ceh-17(np1)*, GR1307 *daf-16(mgDf50)*, and CF1038 *daf-16(mu86)* strains were used. *C. elegans* were grown on NGM agar plates with *E. coli* OP50 bacterial food at 20°C (Brenner, 1974). Animals were obtained by selecting L4 larval stage animals; after 24 hours, animals were used in assays as day 1 adult animals. For aging assays, 5-fluoro-2’-deoxyuridine (FUDR) was not used to suppress progeny production. Animals were gently serially passaged to avoid overcrowding of plates with progeny.

### Microfluidic chip preparation

PDMS microfluidic chips with 24 wells (1.6 mm wide, 0.07 mm deep, 0.4 mm gap between wells) organized in 4 rows and 6 columns were created using a custom mold. A Sylgard 184 silicone elastomer kit (Dow Chemical) was used to make PDMS, which was poured onto the mold to a thickness of 4 mm. Freshly poured PDMS was degassed in a desiccator using a vacuum until air bubbles had dissipated, then placed in a 55°C oven for 18 hours to cure. PDMS was removed from the mold and cut into chips using a razor. These chips were then soaked in *E. coli* OP50 culture overnight, washed with water and ethanol, and left to dry for a week to decrease hydrophobicity before use in swimming assays.

### Kanamycin-treated E. coli OP50 preparation

Kanamycin-treated *E. coli* OP50 food solution was prepared as previously described (Huang, Singh, & Hart, 2017). In brief, *E. coli* OP50 was streaked onto LB agar plates and cultured overnight at 37°C. A single colony was used to inoculate 100 mL of liquid LB and cultured at 30°C shaking at 220 rpm for approximately 12 hours. The culture was grown until it reached an optical density of 2-2.5 at 550 nm and concentrated to a final OD550 of 10 (OD determinations made using diluted cultures to stay within the linear range of the spectrophotometer). 0.2 mg/mL Kanamycin was added and the culture was placed at 4°C for one week to yield a static bacterial culture. Kanamycin-treated OP50 was discarded after 6 weeks of antibiotic treatment. Immediately prior to assays, 200 microliters of kanamycin-treated OP50 was pelleted and resuspended in 300 microliters of liquid NGM.

### Swimming behavioral assays

Chips were cleaned of dust and debris using laboratory tape. Microfluidic chips were then placed into 35 × 10 mm Petri dishes. Each well was loaded with kanamycin-treated OP50 in liquid NGM until a dome of the liquid droplet was visible, but no dark shadows were visible (approximately 0.5 microliters). Water was added to the Petri dish until just level with the chip surface and paraffin oil was layered over water to prevent evaporation. Before loading in individual wells, animals were picked to an unseeded plate to avoid contaminating wells with additional food. In assays with multiple genotypes, loading order always changed between trials. Animals were recorded swimming at 30 frames per second for six hours with a Grasshopper3 4.1MP Mono USB3 Vision camera (GS3-U3-41C6M) mounted on a Zeiss Discovery V20 microscope with a 1.25x objective providing 14.8x magnification. For image capture, we used FlyCap2 version 2.12.3.2 or SpinView version 1.13.0.33. Image resolution was 2048 × 1600, 270 × 270 per chamber with approximately 1,800 pixels per animal. Representative video available at DOI: 10.5281/zenodo.3604455.

For pumping analysis, videos were recorded using the same setup but at 72.0x magnification to allow for visualization of the pharynx. For quiescent bout analysis, these videos were scaled to match the resolution of all other videos to ensure that quiescent bouts were called in a similar manner. After identification of quiescent bouts, the original videos were manually analyzed for pumping status. If the pharynx was for any reason not visible during a bout (self-occlusion or debris), those bouts were not counted (5 instances).

Beat rate was manually collected by analyzing the first minute of swimming. One beat was defined as a full-body bend by the animal in both directions. The experimenter was blinded during this analysis.

### Automated well detection

As a pre-processing stage, wells were automatically segmented using MATLAB’s connected component algorithm (function ***bwboundaries***) on the output of an edge detector (Kimia, Li, Guo, & Tamrakar, 2018). The top 24 connected components were then selected – each corresponding to a different well.

### Activity level analysis

The behavior of each animal was analyzed for each well independently. We implemented a simple ‘active’ vs. ‘quiescent’ classifier by considering the binary output of an edge detector (Kimia *et al*., 2018) and computing the edge difference between consecutive frames using the Jacquard index defined as: *IoU = Area of Overlap / Area of Union*, where *Area of Overlap* is the number of edge pixels present in two consecutive frames and *Area of Union* is the sum of the total number of edge pixels across consecutive frames. If the *IoU* was above a threshold θ = 0.9, the behavior was set to quiescent. Otherwise, the behavior was set to active. Using edges rather than pixels yielded robustness to noise compared to simpler systems based on pixels (Restif *et al*., 2014). A final class label was computed by voting between the two behaviors over a 1 second (30 frames) time window. The activity threshold θ was treated as a hyperparameter and a grid search strategy was employed to identify the optimal value. By systematically varying θ from 0.1 to 1 (with a step size of 0.05) and computing the Jacquard index between the estimations and manually-annotated labels, we were able to automatically determine optimal θ* that yielded the highest agreement. The annotated dataset was comprised of active/quiescent labels from video recordings of 8 held out animals, each video lasting 20 minutes at 30 frames per second. The optimal threshold value θ* was identified to be 0.9, and yielded an agreement of 92%.

### Analysis of binary quiescence data

For each hour, binary quiescence data for each animal were analyzed using MATLAB to determine fractional quiescence, time to first bout, bout number, and bout duration. First, data for each animal was iterated through to determine the start and end times for every quiescent bout across the six-hour assay. Arbitrarily, we defined the minimum duration of a quiescent bout as three seconds. Fractional quiescence for each hour was determined by summing the duration of quiescent bouts observed that hour and dividing this by time in that hour. To determine bout number per hour, the number of bouts observed in an hour were tallied. Bout duration was determined by dividing the total time spent in quiescent bouts by the number of bouts that occurred in that hour. For bout number and duration, if a bout crossed over multiple hours, it was counted for each hour. For fractional quiescence, if a bout crossed over multiple hours, only the portion of the bout that occurred in a given hour contributed to the fractional quiescence for that hour. The time to first bout was calculated by determining the second in which the first quiescent bout for an animal occurred.

### Kinematic analysis

The approach is based on a pose tracking algorithm developed in-house (Guo *et al*., 2018). Briefly, nine body landmarks were chosen along the medial axis of the animal body from head to tail. We collected ten representative video sequences (each corresponding to a different animal) with a total length of 1,200 frames (30 frames per second). Ground-truth poses were manually annotated every 10 frames by marking the location of the nine body landmarks. To train and test our computer vision system, we used a leave-one-video out procedure. Detailed evaluation can be found in (Guo *et al*., 2018). On average, 83% of the landmarks were correctly detected (*i*.*e*., within a 5px radius around the ground-truth landmark). This strategy outperformed competing systems for human pose estimation, specifically the original deformable part model (Yang & Ramanan, 2013) and a leading neural network architecture for human pose estimation, the convolutional pose machine (Wei, Ramakrishna, Kanade, & Sheikh, 2016). These positional landmarks were then used to compute metrics informative of body movements, as defined by (Restif *et al*., 2014). Since these measures rely heavily on canonical reorientation of the worm with respect to its head, differentiating the head from the tail becomes particularly important. To alleviate this concern, we specifically ran a second round of tracking, using the Hungarian algorithm, manually supervised by one-click per video by a trained expert. Representative video of tracking available at DOI: 10.5281/zenodo.3606717.

### Statistical analysis

Statistical analysis was performed using GraphPad Prism 7.0 software (La Jolla, CA). Statistical significance in fractional quiescence, bout number, and bout duration was determined using a 2-way ANOVA and Dunnett’s multiple comparisons testing. Statistical significance for time to first quiescent bout was determined using Kruskal-Wallis and Dunn’s multiple-comparison tests. A value of p < 0.05 was used to determine statistical significance (* p < 0.05, ** p < 0.01, *** p < 0.001). Error bars in figures represent the standard error of the mean.

## Results

### Behavioral analysis of swimming C. elegans

Analysis of *C. elegans* movements during swimming is critical for the analysis of diverse circuits and behavior. To further characterize *C. elegans* locomotion in liquid media, we took advantage of a new computer-vision system for analysis of swimming animals briefly described by (Guo *et al*., 2018) and assessed changes in *C. elegans* locomotory behaviors during prolonged swimming. Individual young adult animals were placed in small liquid drops with static bacteria as food and high-frame-rate, high-resolution videos were obtained over a 6-hour period. For this analysis, we examined both control animals and *egl-4* mutant animals, which are known to have altered locomotion. The cGMP dependent protein kinase encoded by the *egl-4* gene is critical for a wide range of behaviors, including arousal, locomotion, and sleep (Raizen *et al*., 2008). *egl-4(n477)* loss of function animals and *egl-4(ad450)* gain of function animals were expected to show opposing differences in locomotory behaviors.

### Accurate assessment of locomotory behaviors during swimming

To determine if the computer vision program detected changes in locomotion, we analyzed recordings of 12 wild type animals and 12 animals for each of the *egl-4* mutant alleles. We undertook computer vision system analysis for two time intervals within the 6-hour recording window: the first 30 minutes of swimming (early), when quiescent bouts are unlikely to occur were compared to 30 minutes of swimming (late) about 4 hours (228.25 ± 4.28 minutes) into the videos. For ease of comparison to previous work, we used locomotion parameters that were previously defined in the swimming tracking program CeleST (Supplementary Table 1). To undertake analysis for these locomotion parameters, it was necessary to assign the head and tail of each animal. Initial head and tail assignments were made computationally, with manual re-assignment as necessary after curling bouts. The resulting description of locomotion parameters is shown in Figure 1 and yields values comparable with those obtained using CeleST (Restif *et al*., 2014) (Supplementary Table 2).

**Figure 1:**
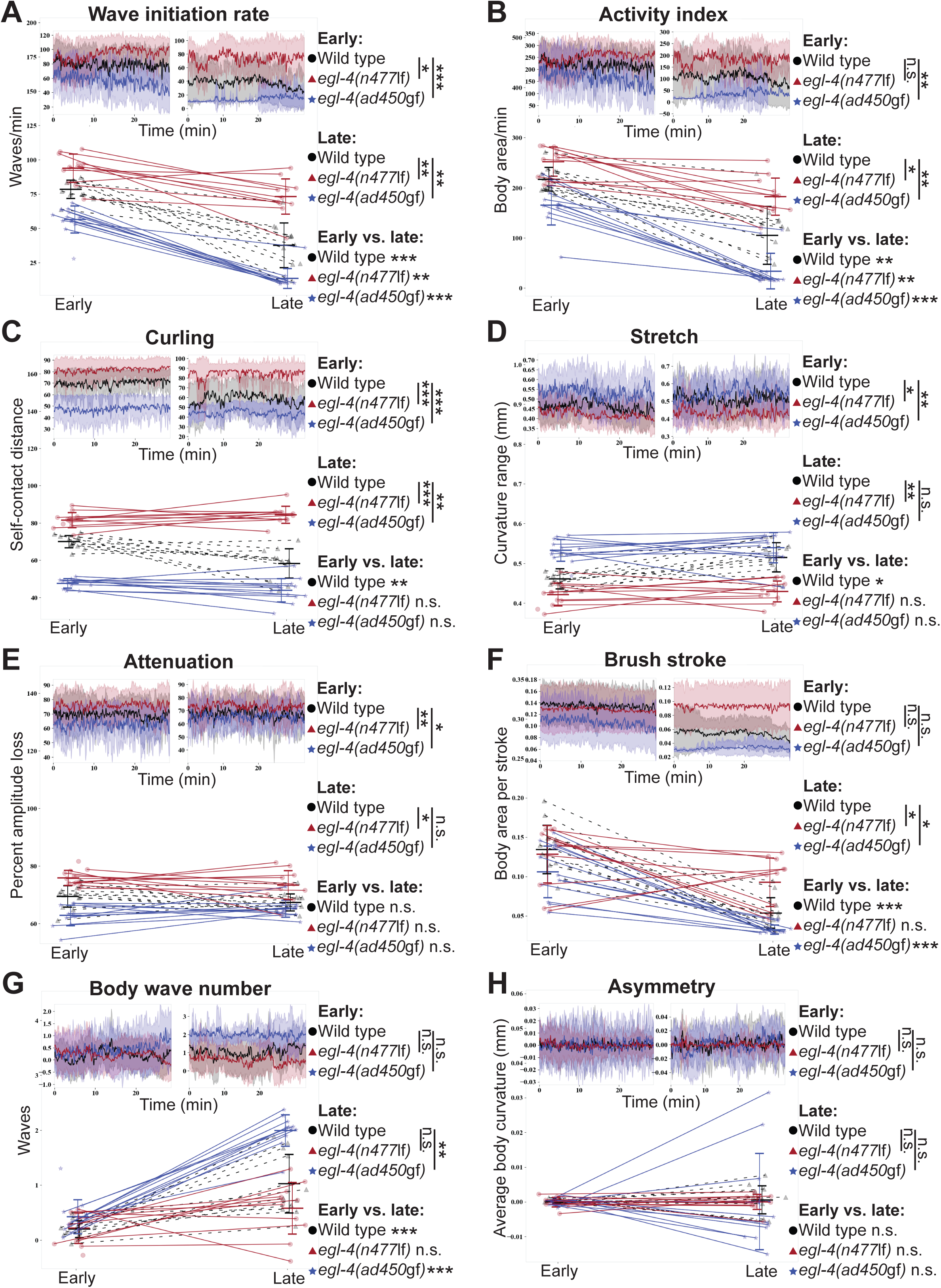
Locomotion analysis of *egl-4* mutant animals during prolonged swimming. Evaluation of parameters (A) Wave Initiation Rate (B) Activity Index (C) Curling (D) Stretch (E) Attenuation (F) Brush Stroke (G) Body Wave Number (H) Asymmetry; each indicative of different aspects of locomotion over two windows of 30 minutes each. For a detailed explanation of these parameters, please refer to (Restif et al., 2014). The ‘early’ time point is at the very beginning of the 6-hour long behavioral assay while the ‘late’ time point is approximately at the 4 hour mark (228.25 ± 4.28 minutes). The choice of these time points was motivated by the ethograms of Figure 2A. Behavioral parameters for 12 animals from each of the three genotypes: wild type (black), *egl-4(n477*lf) (red), and *egl-4(ad450*gf) (blue) are shown here. The within-group mean temporal course of each parameter is shown as insets in the respective panel, with the early time point on the top left and the late time point on the top right. The average parameter value (over the 30 minute window) for each individual animal is shown in the respective panel; corresponding early/late points are connected by dashed/straight lines. The non-parametric Kruskal-Wallis test (with Bonferroni correction for multiple comparisons) was used for testing significance of inter-group differences within and across time points, as well as within-group differences across time points. Error bars indicate ± S.E.M. * P < 0.05; ** P < 0.01; *** P < 0.001.

As expected, the locomotory behavior of *egl-4* animals differed from wild type animals; loss and gain of function animals are expected to have a diametrically opposed impact on behaviors. For example, at both early and late time points, average wave initiation rate and activity index were decreased in *egl-4(ad450*gf) animals and increased in *egl-4(n477*lf) animals, compared to wild type animals (Figure 1A-B). We also found increased curling activity in *egl-4(ad450*gf) animals and decreased curling activity in *egl-4(n477*lf) animals compared to wild type at both early and late time points (Figure 1C). At the early time point, differences in stretch were observed, as curvature range was found to be increased in *egl-4(ad450*gf) animals and decreased in *egl-4(n477*lf) animals when compared to wild type (Figure 1D). Differences from wild type were also observed in attenuation during the early time point, with increased body wave attenuation in *egl-4(ad450*gf) animals and decreased body wave attenuation in *egl-4(n477*lf) animals (Figure 1E). At the later time point, *egl-4(n477*lf) animals showed increased brush stroke and *egl-4(ad450*gf) animals showed decreased brush stroke compared to wild type (Figure 1F). We also observed a significant increase in body wave number of *egl-4(ad450*gf) animals compared to wild type (Figure 1G). *egl-4(n477*lf) animals at the late time point showed increased curvature range and body wave attenuation compared to wild type (Figure 1D-E). The large number of diametrically opposed differences observed in *egl-4* animals suggests that the Guo, *et al* computer vision program can accurately discriminate between normal and mutant locomotion.

### Prolonged swimming changes locomotory behaviors

We compared locomotion parameters across time, comparing early *versus* late time points after prolonged swimming within genotypes. Wave initiation rate and activity index decreased in wild type, *egl-4(n477*lf), and *egl-4(ad450*gf) animals over time (Figure 1A-B). Body wave number increased and brush stroke decreased in wild type and *egl-4(ad450*gf) animals between early and late time points (Figure 1F-G). Finally, curling activity and stretch increased in wild type animals over time (Figure 1C-D). The differences observed in each genotype show that not only can we measure the effects of prolonged exercise on locomotion using the Guo, *et al* system and these parameters, but that changes in locomotion after prolonged swimming differ based on genotype. Overall, the changes observed at the late time point are consistent with less vigorous locomotion after four hours of swimming.

### Swimming behavioral states can be represented using binary data or Hidden Markov models (HMM)

One drawback of detecting locomotion changes using the Guo, *et al* program is that the analysis is computationally intensive and takes a substantial amount of time to run (approximately 10 seconds/frame for pose estimation). Therefore, we focused on quiescent behavior during prolonged swimming and developed an edge-detection program, edgeEIQ, that more efficiently identifies EIQ bouts in swimming animals. Knowing that diminished EGL-4 activity decreases quiescent behavior after prolonged swimming (Ghosh & Emmons, 2008), we used edgeEIQ to detect quiescent bouts on the video used above. Ethograms constructed after analysis of wild type and *egl-4* mutant animals showed clear differences in EIQ bouts for each genotype. The downside of this approach was that the activity threshold was a manually selected parameter (manual optimization of the threshold for behavior to be determined quiescent and EIQ set at minimum duration of 3 seconds). As an alternative, we explored using an unsupervised Hidden Markov Model (HMM) to extract latent underlying state temporal sequences of activity. This circumvented the need for an explicit selection of EIQ minimum duration or other model parameters. Modeling was done using the open-source package ***hmmlearn*** in Python. The two-state HMM yielded a latent state sequence that was qualitatively similar to the ethogram constructed from the manually thresholded edgeEIQ binary data (Figure 2A). In a three-state HMM, the state that seems to correspond best to the inactive state from the manually thresholded binary ethogram seems to be atomic in nature, and thus cannot be further decomposed. The state corresponding best to the active state from the manually thresholded binary ethogram decomposed into two states (Supplementary Figure 1A). We found that HMMs with more than three states resulted in degenerate latent states, *i*.*e*., states for which more than one observation is very rare. We can identify such states from analyzing the transition matrices (Supplementary Figure 1B-C) and locating states for which all outgoing transition probabilities are close to zero.

**Figure 2:**
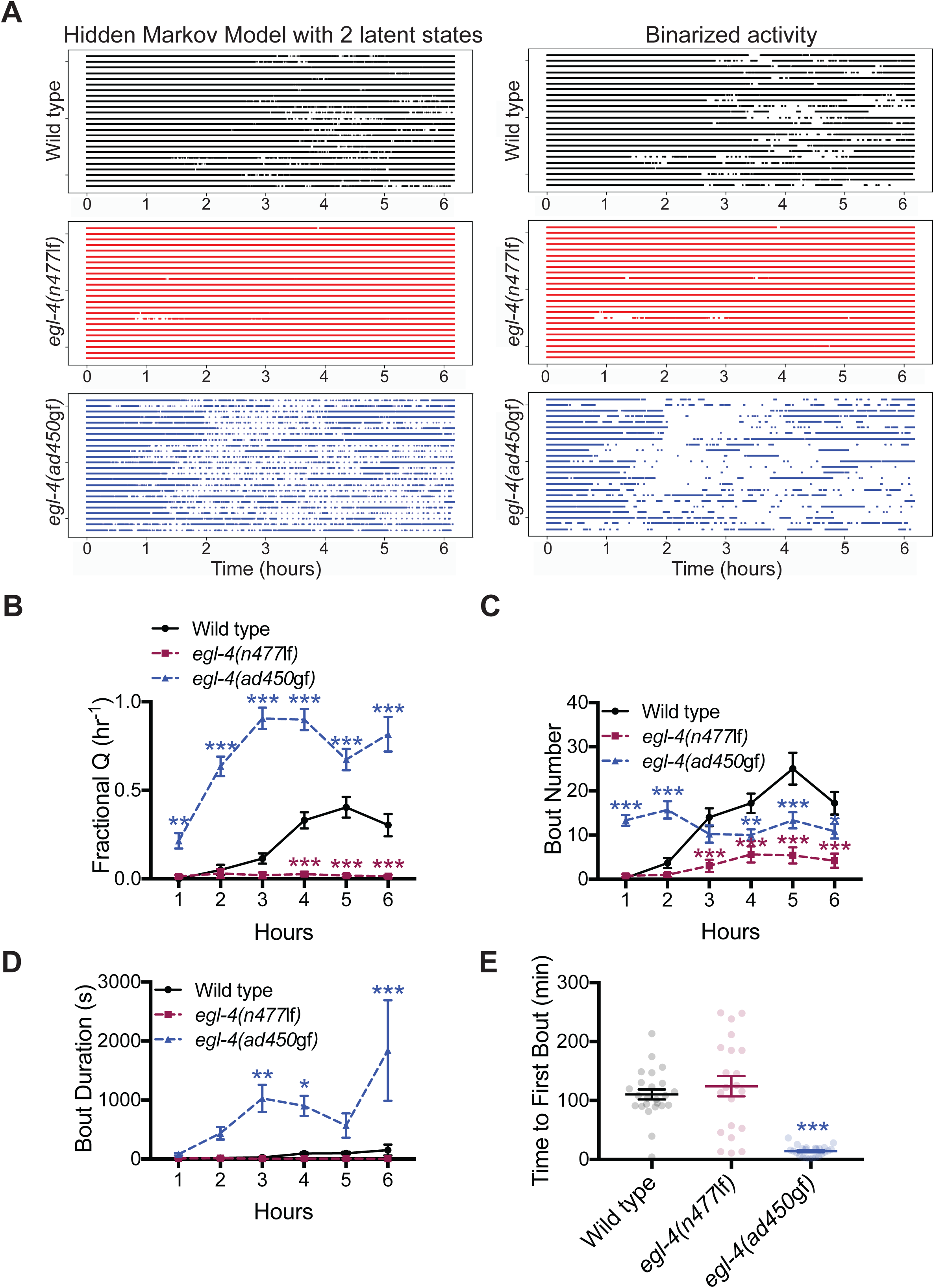
Analysis of exercise-induced quiescence in *egl-4* mutant animals. 24 animals per genotype. (A) Left panel: Ethograms generated using an unsupervised Hidden Markov Model for wild type, *egl-4(n477*lf), and *egl-4(ad450*gf) animals. Each row represents the latent states of a single animal over the course of the 6 hour experiment. Left panel: Ethograms constructed from manually thresholded binary activity data. Filled (or empty) regions can be interpreted as an ‘active’ (or ‘inactive’) state, respectively. (B) On average, the *egl-4(n477*lf) animals showed decreased fractional quiescence (fraction of each hour spent in quiescent bouts) compared to wild type animals during hours 4 through 6. *egl-4(ad450*gf) animals showed increased average fractional quiescence at all time points compared to wild type animals. (C) *egl-4(n477*lf) animals showed decreased average number of bouts (per hour) in hours 3 through 6, compared to wild type animals. *egl-4(ad450*gf) animals showed increased average number of bouts for hours 1 and 2, and decreased average number of bouts in hours 4 through 6, compared to wild type. (D) Average duration of quiescent bouts did not differ between wild type and the *egl-4(n477*lf) animals. *egl-4(ad450*gf) animals showed increased average bout duration during hours 3, 4, and 6. Animals from three independent biological replicates (3 different days). 2-way ANOVA and Dunnett’s multiple comparisons test. (E) *egl-4(ad450*gf) animals initiated quiescent bouts earlier than wild type animals, while *egl-4(n477*lf) animals were not different. Kruskal-Wallis test and Dunn’s multiple comparisons test. Error bars indicate ± S.E.M. * P < 0.05; ** P < 0.01; *** P < 0.001.

To explore transitions between states predicted by HMM modeling, we computed the log-transformed transition count matrices (Supplementary Figure 1B-C). Prior to this, the per-frame latent state sequences were clumped into bouts of length 30-seconds. The equivalent latent state of the bout was assigned as the statistical mode of the latent states of the constituent frames. The complete absence of state 2 in *egl-4(ad450)* gain of function animals, coupled with the relative infrequency of transitions between states 1 and 2 in the wild type animals lends support to the aforementioned statement concerning the atomicity of the latent states, *i*.*e*., whether or not a given state can be decomposed further into unique latent states (Supplementary Figure 1B,C). The most straightforward conclusion from HMM modeling of locomotory behavior is that swimming *C. elegans* have a single inactivity state. Additionally, swimming *C. elegans* likely have two activity states, which is consistent with previous work (McCloskey, Fouad, Churgin, & Fang-Yen, 2017).

### Accurate assessment of quiescent behavior during prolonged swimming

To confirm that our edgeEIQ program could detect previously described differences in EIQ, we compared wild type animals swimming in the presence of food to animals carrying previously described *egl-4* mutant alleles. *egl-4(n477)* loss of function animals were predicted to show decreased EIQ and *egl-4(ad450)* gain of function animals were expected to show increased EIQ. New six-hour videos of swimming animals were recorded and analyzed with the program. The fraction of time quiescent (fractional quiescence) was determined for each hour in individual animals across the six-hour experiment (Figure 2B). At every time point, *egl-4(ad450)* gain of function animals showed increased fractional quiescence compared to wild type, while *egl-4(n477)* loss of function animals showed decreased fractional quiescence for hours four, five, and six, which is consistent with our prediction and previous work (Ghosh & Emmons, 2008). Next, we examined the average number of quiescent bouts in each hour and the average quiescence bout duration for each hour. *egl-4(n477)* animals had decreased bout numbers, starting in hour three and onward (Figure 2C). *egl-4(ad450)* gain of function animals showed increased bout numbers per hour during the first two hours of swimming and decreased bout numbers for hours four through six (Figure 2C) and had increased bout durations at almost all time points, with the exception of hour one (Figure 2D). Finally, we examined when wild type and *egl-4* mutant animals first entered a quiescent bout after prolonged swimming. On average, *egl-4(ad450)* gain of function animals showed quiescence at an earlier time than wild type animals (first quiescent bout of ≥3 seconds, Figure 2E). The loss of *egl-4* function did not alter quiescent bout onset. Overall, these results obtained are entirely consistent with previous work and suggest that the edgeEIQ program can robustly identify differences in EIQ and other quiescent behaviors of swimming animals.

### EIQ does not require pathways necessary for developmentally-timed sleep or stress-induced sleep

A well-characterized quiescent state in *C. elegans* is sleep (Hill, Mansfield, Lopez, Raizen, & Van Buskirk, 2014; Raizen *et al*., 2008). To determine whether sleep was occurring during EIQ bouts, we manually examined quiescent bouts in wild type animals to determine if feeding was occurring. In *C. elegans*, pharyngeal pumping can be used as a metric for food intake and feeding. During each hour, pharyngeal pumping was observed in the majority of quiescent bouts; very few bouts were observed where no pharyngeal pumping occurred (Figure 3A). However, during several quiescent bouts, animals did not pump in the first part of the quiescent bout, then resumed pumping for the remainder of the bout. We called these ‘mixed bouts’ and found that the percentage of mixed bouts increased as quiescent bout duration increased (right panel, Figure 3A). However, in the majority of quiescent bouts animals were pumping, suggesting that most *C. elegans* EIQ bouts are not sleep.

**Figure 3:**
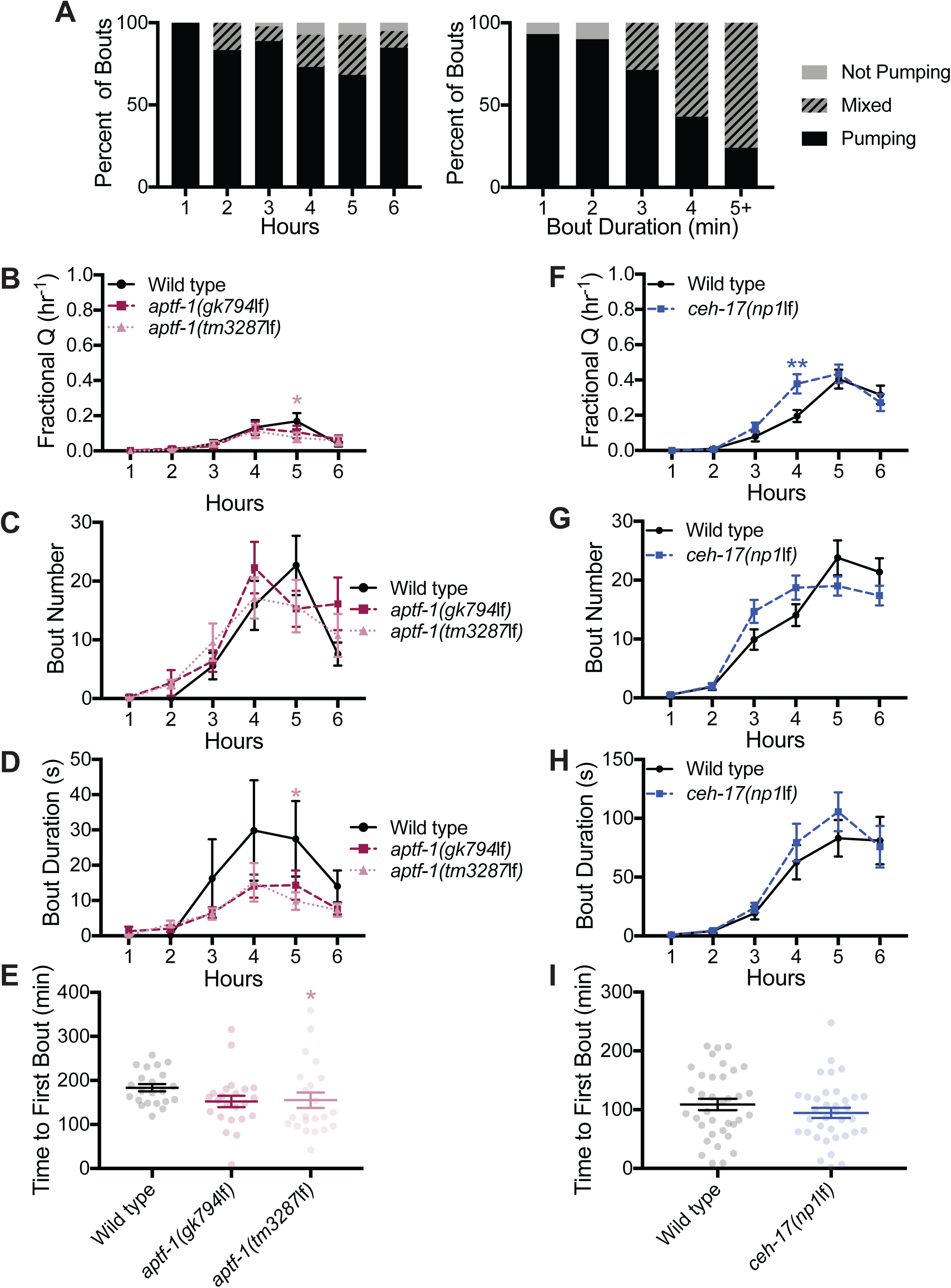
Exercise-induced quiescent bouts are not sleep. (A) Behavior of wild type animals during quiescent bouts was classified as ‘pumping’ (exhibited pharyngeal pumping throughout a quiescent bout), ‘mixed’ (began a quiescent bout without pumping, and resumed pumping midway through the bout), and ‘not pumping’ (no pharyngeal pumping). Left panel: The majority of quiescent bouts were classified as pumping, regardless of when they occurred. Right panel: When classified based on bout duration, most bouts lasting 3 minutes or less were classified as not pumping, while bouts longer than 4 minutes were usually classified as mixed bouts. 199 total quiescent bouts classified drawn from 5 animals. Loss of function *aptf-1(gk794*lf) and *aptf-1(tm3287*lf) animals did not differ from wild type in fractional quiescence (A), bout number (B), or bout duration (C), with the exception of fractional quiescence (B) and bout duration (B) of *aptf-1(tm3287*lf) animals at hour 5. 2-way ANOVA and Dunnett’s multiple comparisons test. (E) Average time to first bout was slightly sooner in *aptf-1(tm3287*lf), but not *aptf-1(gk794*lf), animals *versus* wild type. Kruskal-Wallis test and Dunn’s multiple comparisons test. n = 24 per genotype. Loss of function *ceh-17(np1*lf) showed no difference in fractional quiescence (F), bout number (G), and bout duration (H), with the exception of increased fractional quiescence at hour 5 (F). 2-way ANOVA and Dunnett’s multiple comparisons test. (I) No difference in time to first bout was observed between *ceh-17(np1*lf) animals and wild type. Kruskal-Wallis test and Dunn’s multiple comparisons test. n = 24 per genotype. Error bars indicate ± S.E.M. * P < 0.05; ** P < 0.01; *** P < 0.001.

Examination of mutant strains confirms that there is little mechanistic overlap between EIQ and previously defined *C. elegans* sleep states. Changes in EGL-4 kinase activity impact all known types of *C. elegans* sleep (Hill *et al*., 2014; Raizen *et al*., 2008; You, Kim, Raizen, & Avery, 2008). Perturbation of EGL-4 also alters sensory response and changes locomotion in waking animals (Figure 1A-D). The AP2 transcription factor APTF-1 is specifically required for locomotion quiescence during *C. elegans* developmentally-timed sleep (Turek, Lewandrowski, & Bringmann, 2013) and the paired homeodomain transcription factor CEH-17 is required for locomotion quiescence in stress-induced sleep (Hill *et al*., 2014). We tested *apft-1(gk794)* and *apft-1(tm3287*) loss of function mutants for defects in EIQ and found that loss of APTF-1 does not alter fractional quiescence, bout number, bout duration, or time to first bout, when compared to wild type animals (Figure 3B-E). Although *apft-1(tm3287)* differed from wild type in time to first bout as well as fractional quiescence and bout duration at hour five, similar changes were not observed in *apft-1(gk794)* animals, decreasing confidence that these changes can be attributed to decreased *apft-1* function (Figure 3B,D,E). We found that *ceh-17(np1)* loss of function animals were not different than wild type animals in fractional quiescence, bout number, bout duration, or time to first bout (Figure 3F-I). Combined, these results suggest that different molecular mechanisms underlie EIQ *versus C. elegans* locomotion quiescence during sleep.

### DAF-16/FOXO function delays initiation of quiescent bout cycling after prolonged swimming

The DAF-16/FOXO transcription factor plays a critical role in response to multiple stressors, including sleep restriction (Driver, Lamb, Wyner, & Raizen, 2013; Henderson & Johnson, 2001). We examined both *daf-16(mgDf50)* and *daf-16(mu86)* loss of function mutant animals for changes in EIQ timing. *daf-16(mgDf50)* animals showed increased fractional quiescence at hours three, four, and six (Figure 4A), as well as increased bout number and bout duration at hours three and four (Figure 4B-C). However, these differences were not observed in *daf-16(mu86)* animals. Both loss of function strains showed decreased time to first quiescent bout (Figure 4D). Because this last defect was seen in both mutant *daf-16* strains, we suggest that DAF-16 plays a role in the response to stress caused by prolonged swimming that is important to determining EIQ onset.

**Figure 4:**
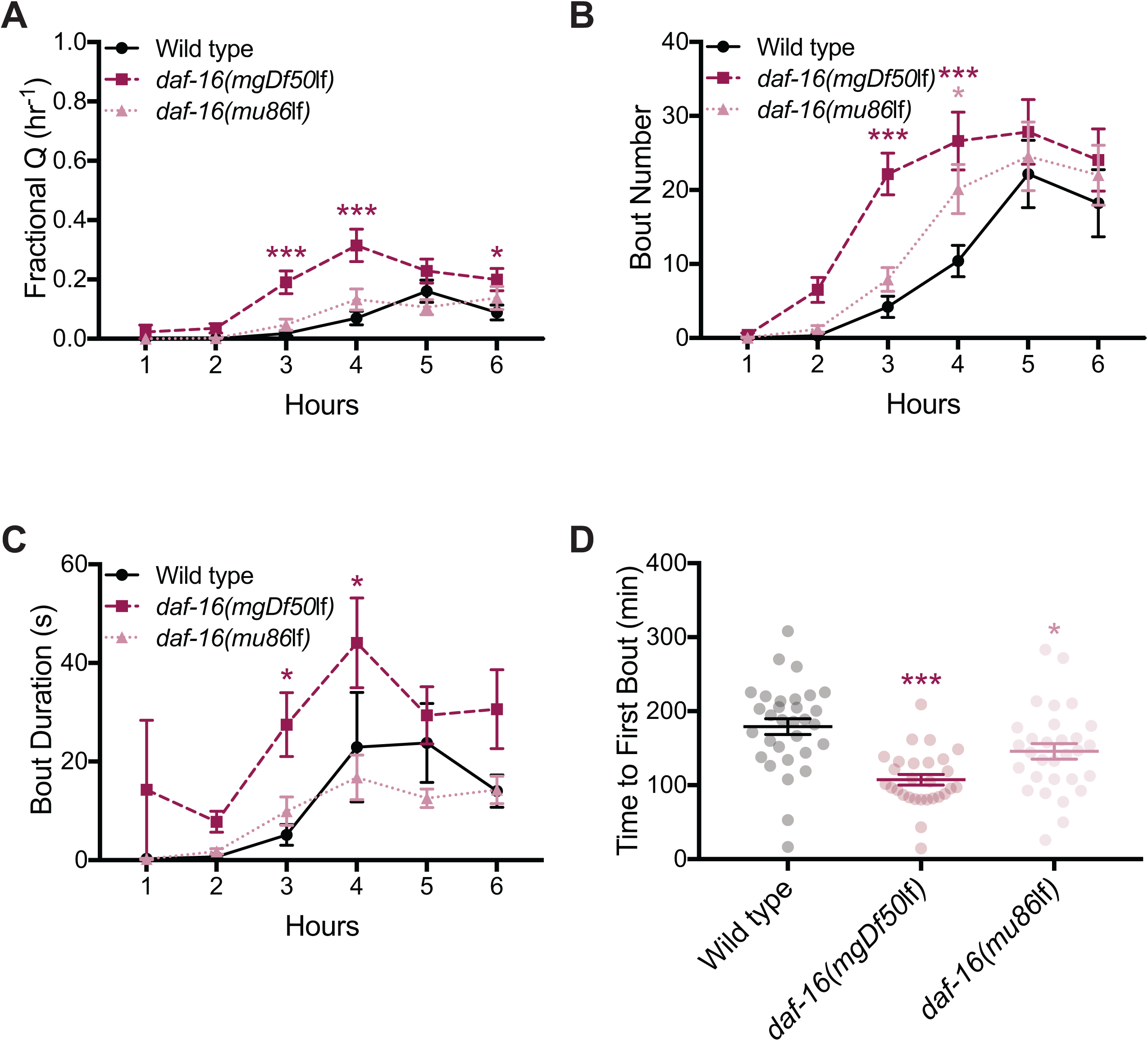
Analysis of exercise-induced quiescence in *daf-16* mutant animals. (A) Loss of function *daf-16(mgDf50*lf) animals showed increased average fractional quiescence, compared to wild type at hours 3, 4, and 6. This difference was not repeated in *daf-16(mu86*lf) animals. (B) *daf-16(mu86*lf) animals showed increased average number of bouts per hour, compared to wild type in hour 4, and *daf-16(mgDf50*lf) animals showed an increase during hours 3 and 4. (C) *daf-16(mgDf50*lf) animals showed increased average bout duration, compared to wild type during hours 3 and 4, while *daf-16(mu86*lf) animals showed no difference. 2-way ANOVA and Dunnett’s multiple comparisons test. (D) The *daf-16(mgDf50*lf) and *daf-16(mu86*lf) animals both showed decreased average time to first bout, compared to wild type. Kruskal-Wallis test and Dunn’s multiple comparisons test. n = 32 per genotype. Error bars indicate ± S.E.M. * P < 0.05; ** P < 0.01; *** P < 0.001.

### Aged animals enter quiescence at an earlier time and show increased quiescence during prolonged swimming

With age, organisms experience loss of muscle mass, known as sarcopenia, which is believed to contribute to increased frailty, decreased muscle strength, and fatigue in aged populations (Marty, Liu, Samuel, Or, & Lane, 2017). Age-related muscle deterioration has previously been observed in *C. elegans* body wall muscle (Herndon *et al*., 2002), and aged *C. elegans* have deficits in various locomotion assays, including swim rate (Mulcahy, Holden-Dye, & O’Connor, 2013; Restif *et al*., 2014). To test whether aged animals show defective EIQ, we aged wild type animals for 1, 2, 3, 4, 7, and 10 days into adulthood. Then, we quantified differences in EIQ timing. Day 7 and 10 adult animals showed increased fractional quiescence compared to day 1 adults, while day 2, 3, and 4 adults were generally indistinguishable from day 1 adults (Figure 5A). No dramatic differences in bout duration were seen between different aged animals. However, bout number was increased from day 7 at hours two, three, and six and increased in day 10 adults at hours one, two, three, five, and six (compared to day 1; Figure 5B-C). On days 3, 4, 7, and 10, animals began cycling between activity and inactivity more quickly than day 1 adults (Figure 5D). A similar decrease in beat rate per minute was also observed at days 2, 3, 4, 7, and 10 (Supplementary Figure 2). These results suggest that locomotory output declines in aged animals, consistent with diminished muscle function in aging animals. Additionally, we noted that different aspects of EIQ metrics decline with age at different rates; time to first EIQ bout decreased with age more rapidly than fractional EIQ increased with age.

**Figure 5:**
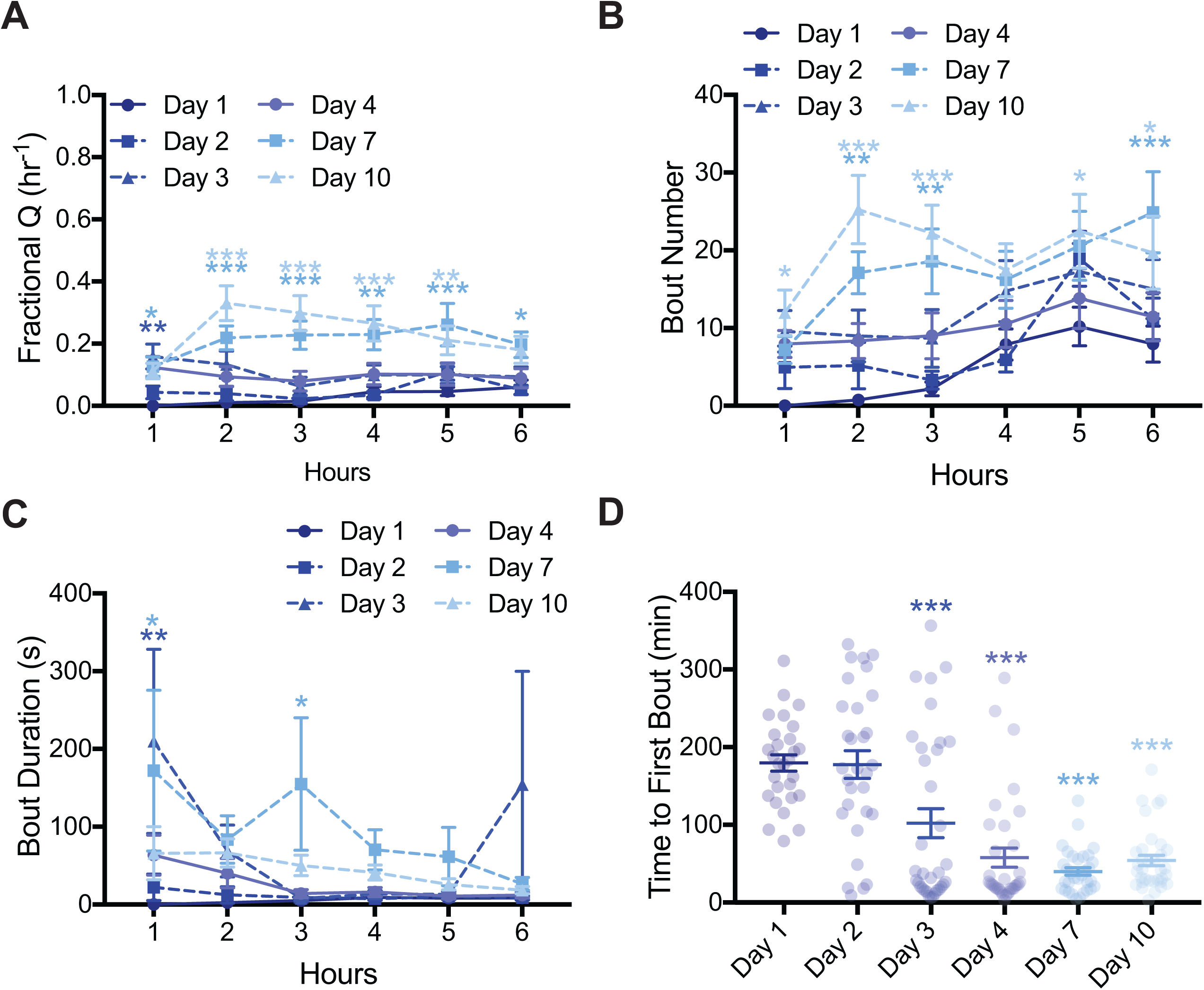
Analysis of exercise-induced quiescence in aged animals. Wild type animals were aged 1, 2, 3, 4, 7, and 10 days into adulthood. (A) Day 3 adult animals showed increased average fractional quiescence, compared to day 1 adult animals at hour 1. Compared to day 1 adult animals, day 7 adult animals showed increased average fractional quiescence at all timepoints, while day 10 animals showed increased average fractional quiescence at hours 2 through 5. (B) An increase in average number of bouts per hour, compared to day 1 adult animals, was observed at day 7 adult animals during hours 2, 3, and 6. Day 10 adult animals also showed increased average number of bouts (per hour) during hours 1, 2, 3, 5, and 6, compared to day 1 adult animals. (C) No difference in average bout duration was found amongst the different age groups, with the exception of increased average bout duration in day 3 adult animals at hour 1 and at day 7 adult animals at hours 1 and 3. 2-way ANOVA and Dunnett’s multiple comparisons test. (D) Day 3, 4, 7, and 10 adult animals all showed decreased average time to first bout, compared to day 1 adult animals. Kruskal-Wallis test and Dunn’s multiple comparisons test. n = 36 per group. Error bars indicate ± S.E.M. * P < 0.05; ** P < 0.01; *** P < 0.001.

## Discussion

Here, we work with computer vision programs that reveal for the first time, in fine detail, changes in *C. elegans* locomotion after prolonged periods of swimming. Increased inactivity was observed after extended swimming, as were differences in swimming locomotory behaviors between wild type and mutant animals. Loss of the EGL-4 cGMP-dependent kinase and DAF-16 FOXO function altered EIQ after prolonged swimming. However, loss of the proteins APTF-1 and CEH-17, which are required for developmentally-timed sleep and stress-induced sleep in *C. elegans*, respectively, did not affect EIQ. Aged animals showed increased EIQ. Based on examining pharyngeal pumping, animals are actively feeding during the majority of EIQ bouts, indicating that EIQ is usually not a sleep state. Computer-vision programs used here enabled in-depth analysis of EIQ and locomotory behaviors across time and revealed previously undescribed defects. This work and development of these automated analysis strategies enables future work that will interrogate the molecular pathways underlying behaviors associated with exercise and fatigue.

*C. elegans* locomotory behavior after prolonged swimming has not been thoroughly studied. Previous studies were limited by reliance on manual annotation, which hinders research depth, or by reliance on constrictive microfluidic devices, which may induce mechanical stress (Ghosh & Emmons, 2008; Gonzales, Zhou, Fan, & Robinson, 2019). Using both of our new systems, we can provide a detailed analysis of how *C. elegans* locomotory behaviors change after prolonged swimming. When comparing wild type and *egl-4* mutant animals, differences were found in multiple parameters, including wave initiation rate at early and late time points. Interestingly, there were also differences in which locomotion parameters changed over time between wild type and mutant animals. For example, curling activity and stretch were found to change over time in wild type animals, but not in *egl-4* mutant animals. In future studies of *C. elegans* fatigue, parameters like wave initiation rate and brush stroke will likely provide information about how vigorously an animal swims and can be used to track decreased muscle output after prolonged swimming exercise.

It is important to note that the computer-vision program developed by Guo, *et al* and used herein for analysis of swimming locomotory behavior is computationally expensive and requires substantial time for processing. To increase efficiency, we developed edgeEIQ which is faster, but less comprehensive, as it only distinguishes active and inactive states during swimming. Loss of function of the gene *egl-4* has previously been associated with decreased quiescence after prolonged swimming (Ghosh & Emmons, 2008, 2010). To test whether edgeEIQ could detect the impact of *egl-4* mutations on EIQ, we analyzed the behavior of gain and loss of function mutants of *egl-4*. As expected, gain and loss of function mutations in *egl-4* were associated with increased and decreased EIQ activity, respectively. cGMP-dependent protein kinase EGL-4 is also known to promote quiescent activity during all known forms of *C. elegans* sleep (Hill *et al*., 2014; Raizen *et al*., 2008). We originally hypothesized that the quiescent behavior after prolonged swimming might also be a sleep state. Usually sleeping animals will stop both feeding and locomotion. But, we found that pharyngeal pumping was usually observed in animals during EIQ bouts, suggesting that EIQ bouts are usually not sleep. We also used genetic strategies to explore the relationship between sleep and EIQ bouts. The RIS neuron is critical for developmentally-timed sleep, and function of the APTF-1 transcription factor is required for RIS-mediated sleep induction (Turek *et al*., 2013). Likewise, the ALA sensory neuron is required for quiescent behavior during stress-induced sleep (Hill *et al*., 2014). CEH-17 loss alters gene expression in the ALA neuron, and loss of this protein leads to shortened ALA axons and inability to enter a sleep state following cellular stress (Hill *et al*., 2014; Pujol, Torregrossa, Ewbank, & Brunet, 2000; Van Buskirk & Sternberg, 2010). We tested animals lacking *aptf-1* and *ceh-17* function for defects in EIQ to determine whether exercise-induced locomotion quiescence was mediated by pathways involved in mediating locomotion quiescence in developmentally-timed or stress-induced sleep. We found that these mutant animals had normal locomotion quiescence. Therefore, EIQ and sleep are not identical and are likely mediated by overlapping, but distinct molecular and cellular pathways.

Prolonged exercise is stressful. In *C. elegans*, the transcription factor DAF-16/FOXO localizes to the nucleus during cellular stress, where it upregulates genes involved with stress response, including oxidative stress response genes (Henderson & Johnson, 2001; Murphy *et al*., 2003). As prolonged swimming by *C. elegans* leads to transcriptional changes in oxidative stress response genes (Laranjeiro *et al*., 2017), we reasoned that DAF-16 might play a role in EIQ and stress response. Loss of function *daf-16* mutant animals initiated EIQ earlier than wild type animals, suggesting that DAF-16 normally promotes expression of genes that allow animals to endure the stress of exercise for a longer initial swim period of time.

As they age, animals experience deterioration that can result in fatigue, weakness, and sarcopenia. We examined EIQ as *C. elegans* age and, as predicted, older animals showed increased EIQ. Surprisingly, a dramatic decrease in initial swim period time was seen in animals 3 or more days into adulthood. This deterioration is not mirrored in fractional EIQ, as animals only show an increase in fractional EIQ at 7 or more days into adulthood. Further analysis of EIQ should provide insight into the effects of the aging process on muscle function over time. Changes in EIQ in aged *C. elegans* are likely a marker of healthspan and the mechanisms underlying these changes may be conserved mechanisms relevant to fatigue in aging humans.

Here, we developed systems for automatic analysis of *C. elegans* swimming locomotory behavior with a long-range goal of understanding locomotion quiescience and associated mechanisms. The cellular and molecular pathways that communicate fatigue from muscles to the nervous system during exercise remain obscure. In complex animals, afferent neuronal pathways are thought to carry information from the periphery to the central nervous system, which eventually results in a ‘feeling of exhaustion’ that results in decreased locomotion. Energy depletion in peripheral organs could also result in decreased locomotion. Results presented here and other works (Beron *et al*., 2015; Lesanpezeshki *et al*., 2019; Rahman *et al*., 2018) have demonstrated that invertebrates show aspects of fatigue, even though they lack extensive afferent neuronal pathways. Further dissection of these behaviors in invertebrates should provide insight into conserved cellular and molecular pathways involved in fatigue, as well as other aspects of endurance and exercise.

## Conclusions

Computer-vision systems have been developed for accurate analysis of *C. elegans* locomotory behavior during prolonged swimming exercise (over 6 hours). This system is complemented by a faster edge-detection-based system that can detect pauses in locomotion. We find that most prolonged swimming results in exercise-induced quiescence (EIQ) bouts that are dependent on EGL-4/PKG function, but not on the function of genes that are specifically required for *C. elegans* sleep. The timing of EIQ bout initiation is dependent on DAF-16/FOXO function. As *C. elegans* age, the timing, duration, and frequency of EIQ bouts change, consistent with diminished healthspan.

## Acknowledgments

This research was supported by funds from Brown’s Office for the Vice-President for Research (A.C.H. and T.S.). Additional support provided by the Carney Institute for Brain Science, the Center for Vision Research (CVR) and the Center for Computation and Visualization (CCV) at Brown University and a Karen T. Romer Undergraduate Teaching and Research Awards (S.K.). We acknowledge the Cloud TPU hardware resources that Google made available via the TensorFlow Research Cloud (TFRC) program. Some strains were provided by the CGC, which is funded by the NIH Office of Research Infrastructure Programs (P40 OD010440).

---

Supplementary Figure 1: Additional state-space analysis of exercise-induced quiescence in *egl-4* mutant animals. (A) Ethograms generated using an unsupervised Hidden Markov Model for wild type, *egl-4(n477*lf), and *egl-4(ad450*gf) animals similar to Figure 2A. The HMM here was constructed with three latent states. The coloring scheme from Figure 2A is maintained with the only difference being that the new latent state is shown in green. Log-transformed bout transition count matrices are shown for the two state HMM (subpanel B) and the three state HMM (subpanel C).

Supplementary Figure 2: Swimming beat rate in aged animals. Swimming beat rate was manually observed in wild type animals that were aged 1, 2, 3, 4, 7, and 10 days into adulthood. A decrease in beat rate per minute compared to day 1 adult animals was observed in all ages. Kruskal-Wallis test and Dunn’s multiple comparisons test. n = 36 per group. Error bars indicate ± S.E.M. * P < 0.05; ** P < 0.01; *** P < 0.001.

Supplementary Table 1: Swimming behavior parameters, adapted from Restif (2014). See Restif (2014) for equations.

Supplementary Table 2: Swimming behavior parameter comparison between our implementation (Guo, *et al* 2018) *versus* CeleST (Restif *et al*, 2014). As a verification step, a total of n=9, 33.3 second long video clips from 3 individual animals (one from each wild type, *egl-4(n477*lf), and *egl-4(ad450*gf)), were processed side-by-side using strategies in Guo, *et al* 2018 and CeleST from Restif, *et al*, 2014 to generate swimming behavior parameters. Values for curling reported here represent seconds of curling (i.e. self-contact distance is less than a threshold value) per minute of swimming and not self-contact distance per se. CeleST’s tracking algorithm often failed on videos acquired using our setup (potentially due to a combination of image resolution, background illumination, worm-background contrast and challenging body poses). To keep the focus on ensuring comparable behavior parameters, here we provided CeleST with tracked worm skeletons from Guo, *et al* 2018 to obtain parameter values.

## References

Beron, C., Vidal-Gadea, A. G., Cohn, J., Parikh, A., Hwang, G., & Pierce-Shimomura, J. T. (2015). The burrowing behavior of the nematode Caenorhabditis elegans: a new assay for the study of neuromuscular disorders. Genes, Brain, and Behavior, 14(4), 357–368.

Brenner, S. (1974). The genetics of Caenorhabditis elegans. Genetics, 77(1), 71–94.

Driver, R. J., Lamb, A. L., Wyner, A. J., & Raizen, D. M. (2013). DAF-16/FOXO regulates homeostasis of essential sleep-like behavior during larval transitions in C. elegans. Current Biology: CB, 23(6), 501–506.

Ghosh, R., & Emmons, S. W. (2008). Episodic swimming behavior in the nematode C. elegans. The Journal of Experimental Biology, 211(Pt 23), 3703–3711.

Ghosh, R., & Emmons, S. W. (2010). Calcineurin and protein kinase G regulate C. elegans behavioral quiescence during locomotion in liquid. BMC Genetics, 11, 7.

Gomez-Marin, A., Partoune, N., Stephens, G. J., Louis, M., & Brembs, B. (2012). Automated tracking of animal posture and movement during exploration and sensory orientation behaviors. PloS One, 7(8), e41642.

Gonzales, D. L., Zhou, J., Fan, B., & Robinson, J. T. (2019). A microfluidic-induced C. elegans sleep state. Nature Communications, 10(1), 5035.

Guo, Y., Govindarajan, L., Kimia, B., & Serre, T. (2018). Robust pose tracking with a joint model of appearance and shape (No. Arxiv).

Henderson, S. T., & Johnson, T. E. (2001). daf-16 integrates developmental and environmental inputs to mediate aging in the nematode Caenorhabditis elegans. Current Biology: CB, 11(24), 1975–1980.

Herndon, L. A., Schmeissner, P. J., Dudaronek, J. M., Brown, P. A., Listner, K. M., Sakano, Y., … Driscoll, M. (2002). Stochastic and genetic factors influence tissue-specific decline in ageing C. elegans. Nature, 419(6909), 808–814.

Hill, A. J., Mansfield, R., Lopez, J. M. N. G., Raizen, D. M., & Van Buskirk, C. (2014). Cellular stress induces a protective sleep-like state in C. elegans. Current Biology: CB, 24(20), 2399–2405.

Huang, H., Singh, K., & Hart, A. C. (2017). MeasuringCaenorhabditis elegansSleep During the Transition to Adulthood Using a Microfluidics-based System. Bio-Protocol, 7(6). https://doi.org/10.21769/BioProtoc.2174

Jung, S.-K., Aleman-Meza, B., Riepe, C., & Zhong, W. (2014). QuantWorm: a comprehensive software package for Caenorhabditis elegans phenotypic assays. PloS One, 9(1), e84830.

Kimia, B. B., Li, X., Guo, Y., & Tamrakar, A. (2018). Differential Geometry in Edge Detection: accurate estimation of position, orientation and curvature. IEEE Transactions on Pattern Analysis and Machine Intelligence, 1–1.

Laranjeiro, R., Harinath, G., Burke, D., Braeckman, B. P., & Driscoll, M. (2017). Single swim sessions in C. elegans induce key features of mammalian exercise. BMC Biology, 15(1), 30.

Lesanpezeshki, L., Hewitt, J. E., Laranjeiro, R., Antebi, A., Driscoll, M., Szewczyk, N. J., … Vanapalli, S. A. (2019). Pluronic gel-based burrowing assay for rapid assessment of neuromuscular health in C. elegans. Scientific Reports, 9(1), 15246.

Marty, E., Liu, Y., Samuel, A., Or, O., & Lane, J. (2017). A review of sarcopenia: Enhancing awareness of an increasingly prevalent disease. Bone, 105, 276–286.

McCloskey, R. J., Fouad, A. D., Churgin, M. A., & Fang-Yen, C. (2017). Food responsiveness regulates episodic behavioral states inCaenorhabditis elegans. Journal of Neurophysiology, 117(5), 1911–1934.

Mulcahy, B., Holden-Dye, L., & O’Connor, V. (2013). Pharmacological assays reveal age-related changes in synaptic transmission at the Caenorhabditis elegans neuromuscular junction that are modified by reduced insulin signalling. The Journal of Experimental Biology, 216(Pt 3), 492–501.

Murphy, C. T., McCarroll, S. A., Bargmann, C. I., Fraser, A., Kamath, R. S., Ahringer, J., … Kenyon, C. (2003). Genes that act downstream of DAF-16 to influence the lifespan of Caenorhabditis elegans. Nature, 424(6946), 277–283.

Patel, T. P., Gullotti, D. M., Hernandez, P., O’Brien, W. T., Capehart, B. P., Morrison, B., … Meaney, D. F. (2014). An open-source toolbox for automated phenotyping of mice in behavioral tasks. Frontiers in Behavioral Neuroscience, 8, 349.

Pujol, N., Torregrossa, P., Ewbank, J. J., & Brunet, J. F. (2000). The homeodomain protein CePHOX2/CEH-17 controls antero-posterior axonal growth in C. elegans. Development, 127(15), 3361–3371.

Rahman, M., Hewitt, J. E., Van-Bussel, F., Edwards, H., Blawzdziewicz, J., Szewczyk, N. J., … Vanapalli, S. A. (2018). NemaFlex: a microfluidics-based technology for standardized measurement of muscular strength of C. elegans. Lab on a Chip, 18(15), 2187–2201.

Raizen, D. M., Zimmerman, J. E., Maycock, M. H., Ta, U. D., You, Y.-J., Sundaram, M. V., & Pack, A. I. (2008). Lethargus is a Caenorhabditis elegans sleep-like state. Nature, 451(7178), 569–572.

Restif, C., Ibáñez-Ventoso, C., Vora, M. M., Guo, S., Metaxas, D., & Driscoll, M. (2014). CeleST: computer vision software for quantitative analysis of C. elegans swim behavior reveals novel features of locomotion. PLoS Computational Biology, 10(7), e1003702.

Stephens, G. J., Johnson-Kerner, B., Bialek, W., & Ryu, W. S. (2008). Dimensionality and dynamics in the behavior of C. elegans. PLoS Computational Biology, 4(4), e1000028.

Turek, M., Lewandrowski, I., & Bringmann, H. (2013). An AP2 transcription factor is required for a sleep-active neuron to induce sleep-like quiescence in C. elegans. Current Biology: CB, 23(22), 2215–2223.

Van Buskirk, C., & Sternberg, P. W. (2010). Paired and LIM class homeodomain proteins coordinate differentiation of the C. elegans ALA neuron. Development, 137(12), 2065–2074.

Wei, S.-E., Ramakrishna, V., Kanade, T., & Sheikh, Y. (2016). Convolutional Pose Machines. IEEE Computer Vision and Pattern Recognition Conference. Retrieved from http://arxiv.org/abs/1602.00134

Yang, Y., & Ramanan, D. (2013). Articulated human detection with flexible mixtures of parts. IEEE Transactions on Pattern Analysis and Machine Intelligence, 35(12), 2878–2890.

You, Y.-J., Kim, J., Raizen, D. M., & Avery, L. (2008). Insulin, cGMP, and TGF-beta signals regulate food intake and quiescence in C. elegans: a model for satiety. Cell Metabolism, 7(3), 249–257.

