## Supplementary Table 1 for "Discriminating between sleep and exercise-induced fatigue using computer vision and behavioral genetics"

| **Parameter** | **Description** |
| --- | --- |
| Wave initiation rate | Body waves per minute (similar to manually recorded beat rates) |
| Body wave number | Number of waves present in the body at one time |
| Asymmetry | How symmetrical strokes are in each direction |
| Curling | Whether animals are into a circular shape, measured as the distance between the head and the tail |
| Stretch | Measures how deep body bends are |
| Attenuation | Measures how well the amplitude of a wave is maintained down the body |
| Activity index | Normalization of brush stroke, shows how vigorously an animal is swimming |
| Brush stroke | Area that an animal covers in one stroke |
