## Supplementary Table 2 for "Discriminating between sleep and exercise-induced fatigue using computer vision and behavioral genetics"

| **Parameter** | **Measure from CeleST software** | **Measure from our implementation** |
| --- | --- | --- |
| Wave initiation rate | 109.81 ± 17.32 | 89.19 ± 17.82 |
| Body wave number | 0.43 ± 0.07 | 0.33 ± 0.40 |
| Asymmetry | -9.24e-05 ± 0.01 | -0.003 ± 0.002 |
| Curling | 1.27 ± 1.40 | 2.14 ± 2.19 |
| Stretch | 0.57 ± 0.06 | 0.48 ± 0.04 |
| Attenuation | 39.89 ± 24.16 | 69.95 ± 6.37 |
| Activity index | 234.11 ± 41.001 | 252.36 ± 53.75 |
| Brush stroke | 0.096 ± 0.04 | 0.15 ± 0.01 |
