## Supplementary figures and images for "Discriminating between sleep and exercise-induced fatigue using computer vision and behavioral genetics"

### Supplementary Figure 1

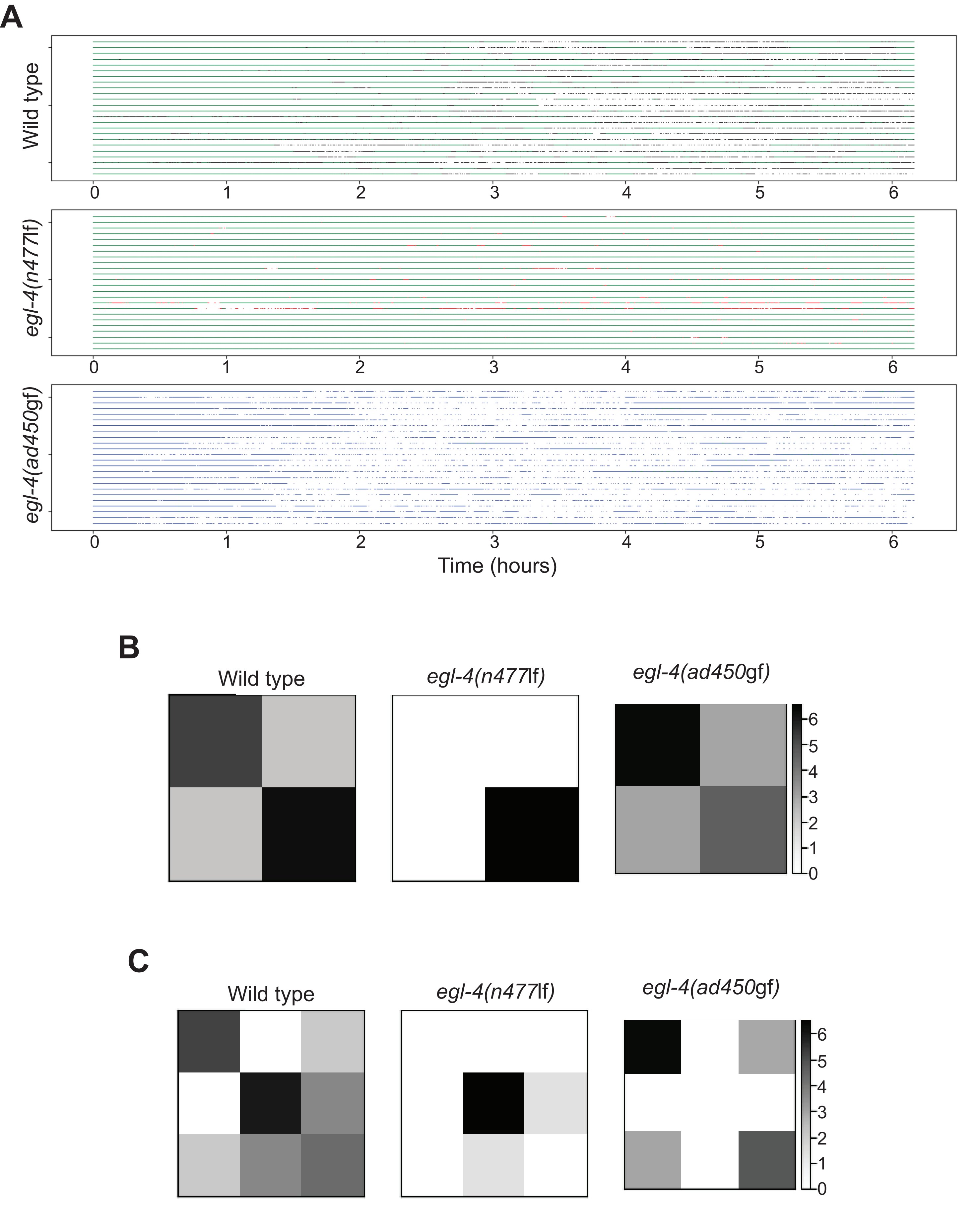

### Supplementary Figure 2

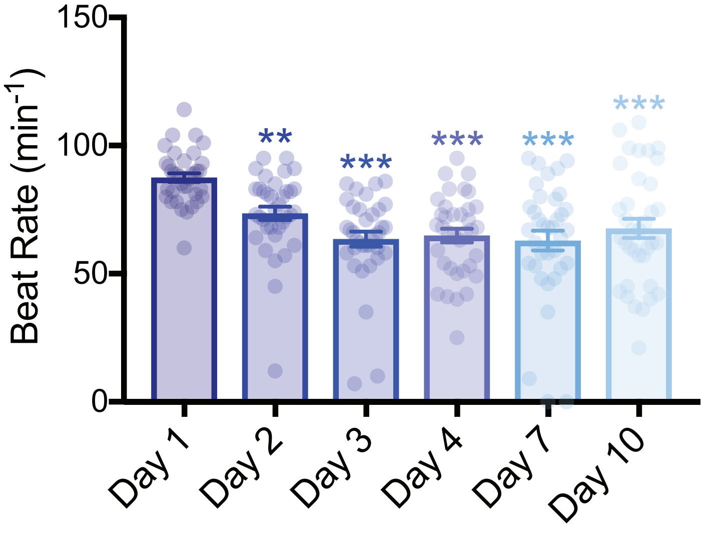
